# AAV9-mediated targeting of natural antisense transcript as a novel treatment for Dravet Syndrome

**DOI:** 10.1101/2024.09.27.615323

**Authors:** Juan Antinao Diaz, Ellie Chilcott, Amanda Almacellas Barbanoj, Anna Keegan, Amy McTague, J Helen Cross, Stephanie Schorge, Gabriele Lignani, Simon Nicholas Waddington, Rajvinder Karda

## Abstract

Dravet syndrome (DS) is a severe childhood onset developmental and epileptic encephalopathy which leads to life-long disability. Symptoms usually manifest in the first year of life and include prolonged severe seizures, developmental delay and severe intellectual disability. DS patients have an increased mortality rate, including sudden unexpected death in epilepsy (SUDEP). Approximately 90% of patients carry a heterozygous loss-of-function mutation in the *SCN1A* gene, which encodes a voltage-gated sodium ion channel, Na_V_1.1. The Na_V_1.1 channel is expressed in the brain and at a lower level, in the heart. Previous studies have identified a long non-coding RNA (lncRNA) which specifically downregulates *SCN1A* gene expression. This natural antisense transcript (NAT) can be modulated by AntagoNATs, small synthetic oligonucleotides developed to inhibit NAT function. In a DS mouse model, AntagoNATs were shown to modulate *Scn1a* expression by targeting the *Scn1a* NAT, improving seizure frequency after repeated administration. Here, we have developed novel AntagoNATs and incorporated these into a clinically relevant adeno-associated virus serotype 9 (AAV9) gene therapy vector, to test in a DS mouse model (*Scn1a*^*+/-*^) and provide a one-off treatment approach.

Eighteen AntagoNATs were tested *in vitro*; from the best performing candidates, we selected two AntagoNAT sequences (K & H) for *in vivo* testing as they had the highest homology (90%) to human *SCN1A* NAT. We administered both vectors to newborn *Scn1a*^*+/-*^ mice via intracerebroventricular (ICV) and intravenous (IV) injection to target the brain and heart. AAV9-AntagoNAT-H significantly increased survival, decreased febrile seizures and reduced spontaneous seizure frequency compared to the PBS control group. When administered at P14 by ICV and IV injection, AAV9-AntagoNAT-H increased survival. In this proof-of-concept study, we have demonstrated for the first time the delivery of AntagoNAT technology via an AAV9 vector and thus offering the possibility of a one-time treatment for DS patients.

## Introduction

Dravet syndrome (DS) is a severe early onset genetic epilepsy, which manifests in the first year of life, with an incidence of 1 in 15,400 to 1 in 40,900 live births world-wide.^1,2^ DS patients exhibit frequent prolonged febrile and afebrile seizures, generalised tonic or hemiclonic seizures, status epilepticus events, with cognitive decline, developmental delay, ataxia and many other comorbidities.^3–5^ Unfortunately, DS patients have an increased mortality rate, including sudden unexpected death in epilepsy (SUDEP).^5^

More than 90% of DS patients harbour a heterozygous loss-of-function mutation in the *SCN1A* gene, which encodes the α subunit of the voltage-gated ion channel, Na_V_1.1.^1^ In the human brain, Na_V_1.1 expression has been identified in the hippocampus, dentate gyrus, CA3 and CA2, layer V/VI of the cortex, granular layer, molecular layer, deep nuclei and Purkinje cells of the cerebellum.^6^ Animal studies have also identified Na_V_1.1 expression in the hippocampus,^7^ thalamus,^8^ deep cerebellar nuclei,^9^ and the spinal cord,^10^ specifically in inhibitory interneurons,^7^ Purkinje neurons^11^ and CA1 pyramidal cells.^12^ Na_V_1.1 has been detected in the human heart and studies have investigated altered electrical cardiac function in DS patients and mouse model, suggesting a possible contribution to SUDEP.^13–16^

Multiple models of DS have been established to study disease mechanisms, which have demonstrated spontaneous seizures, febrile seizures, SUDEP, cognitive deficits, and other comorbidities.^1^ Many of these models have shown a notable dependence on genetic background.^21^ Yu et al.^7^ developed a heterozygous loss of function DS model (*Scn1a*^*+/-*^) where *Scn1a* exon 25 was disrupted. The study revealed, *Scn1a*^*+/-*^ mice on a C57BL/6J strain resulted in a more severe disease phenotype than on a 129/SvJ background. They also developed a mix DS model 129/SvJ:C57BL/6J, demonstrating that Na_V_1.1 loss of function in inhibitory interneurons in the hippocampus resulted in spontaneous seizures and increased mortality.^7^ Cheah et al.^22^ provided further evidence of the importance of GABAergic interneurons in DS by selectively deleting *Scn1a* in forebrain GABAergic neurons, which also resulted in tonic-clonic seizures, febrile seizures and premature death. Another *Scn1a*^*+/-*^ model was developed by deleting the first coding exon of *Scn1a* in a mixed 129/SvJ:C57BL/6J background strain. The first generation of *Scn1a*^*+/-*^ mice demonstrated seizures and premature death.^23^ Further studies on this *Scn1a*^*+/-*^ model have shown a reduction in sodium currents from hippocampal inhibitory GABAergic interneurons. Interestingly, they also noted elevated sodium currents in excitatory pyramidal neurons of the hippocampus.^21^ In addition, studies have also demonstrated altered excitatory CA1 pyramidal neuronal networks during early stages of development in the *Scn1a*^*+/-*^ model.^24^ Together, these studies illustrate the involvement of both inhibitory and excitatory neuronal networks in DS.

Other organs such as the heart have also shown involvement in DS. DS patients demonstrate autonomic symptoms,^13^ in some cases resulting in reduced heart rate variability, which may act as a potential biomarker for SUDEP risk.^25^ DS mouse model have also showed an increase of sodium currents in ventricular myocytes resulting in altered electrical cardiac function.^14^

Overall, the DS mouse model studies have demonstrated that the loss of *Scn1a* in multiple regions of the brain and the association of the heart contributes to disease phenotype. Therefore, there is a need to develop a wide-spread genetic therapy approach to restore function of *SCN1A* in multiple neuronal cells and the heart.

Current anti-seizure medications such as valproate, stiripentol and clobazam^26^ are often ineffective in reducing seizure frequency and other comorbidities associated with DS. Therefore, there is a great need for alternative therapies. Gene supplementation therapy using adeno-associated viral (AAV) vectors have demonstrated tremendous clinical success for severe early onset neurological disorders.^27,28^ However, the *SCN1A* gene (6kb), exceeds the packaging capacity of AAV (∼4.7kb).^29^ Therefore, gene supplementation using a single AAV is not feasible for treating *SCN1A* related DS. Gene supplementation approach using an Adenoviral vector has been developed for DS, as it has a larger packaging capacity than AAV.^30^ Numerous other genetic therapies for DS have been developed which aim to upregulate *SCN1A* expression and some of these are in clinical trials. Stoke Therapeutics have developed an antisense oligonucleotide (ASO) therapy, that aims to increase the synthesis of productive *Scn1a* mRNA.^18^ This treatment is currently in Phase 1/2a clinical trials, where the ASO is delivered via repeated intrathecal delivery to DS patients.^31^ Encoded Therapeutics have also developed an AAV9 based therapy to deliver a transcription factor to upregulate *SCN1A* specifically in GABAergic inhibitory interneurons^19^ which is currently in Phase 1/2 clinical trial.^32^

AntagoNATs are small synthetic oligonucleotides that modulate RNA function, in this case by targeting the long non-coding RNA (lncRNA) from the natural antisense transcript (NAT) of *SCN1A*. These AntagoNATs aim to inhibit the NAT function, thus increasing *SCN1A* mRNA and consequently, Na_V_1.1 protein production.^33^ A reduction of seizure phenotype after repeated intrathecal administration of AntagoNATs to 7-week old DS knock-in mice has been demonstrated.^33^ Critically, this reduction in seizure phenotype required repeated intrathecal delivery.

We sought to improve this approach, by designing new AntagoNAT sequences and incorporating them into a clinically relevant AAV vector. This approach provides a one-off treatment to a clinically relevant DS mouse model.^18^ AAV9 serotype was selected for this study as this has been used extensively in many pre-clinical and clinical studies for central nervous system diseases.^19,28^ Furthermore, we aimed to target the brain and the heart with our therapy, as studies have shown Na_V_1.1 to be expressed in these areas.^7,16,21,22,24^

The *in vitro* data revealed a candidate AntagoNAT sequence H, which showed a higher increase in endogenous *Scn1a* expression when compared to previously published AntagoNAT sequence.^33^ A heterozygous *Scn1a*^*+/-*^ DS mouse model which shows spontaneous seizures, febrile seizures and increased mortality^18,21^ was used in this study. AAV9-AntagoNAT-H was administered to new-born *Scn1a*^*+/-*^ DS mice via intracerebroventricular (ICV) and intravenous (IV) delivery. The results revealed a significant increase in survival and significant reduction in febrile seizures and a reduction in average daily seizure frequency. Furthermore AAV9-AntagoNAT-H delivered to older *Scn1a*^*+/-*^ mice by combination therapy led to an increase in survival.

## Materials and methods

### Cloning

The mouse *Scn1a* NAT secondary structure was predicted using Mfold.^34^ We mapped previously published AntagoNAT sequences^33^ to the resulting secondary structure files, and designed eighteen new sequences targeting different locations of the NAT. These sequences were incorporated into an AAV backbone using gBlocks obtained from Integrated DNA Technologies (IDT, Europe) and cloned using InFusion cloning (Takara Bio, Europe). For the *in vitro* study, RNA polymerase II promoter, CMV was placed upstream of reporter gene, enhanced green Fluorescent Protein (GFP), with mir-155 sequences flanking the AntagoNAT sequences to allow stable expression;^35^ AAV-CMV-eGFP-mir-155-AntagoNAT-mir155-WPRE (Fig. 1A). For the *in vivo* study, AntagoNAT sequences were driven by a RNA polymerase III promoter, U6. The construct also contained an additional CMV promoter to allow the expression of reporter gene, eGFP; AAV-U6-AntagoNAT-CMV-eGFP-WPRE (Fig. 1B).

**Figure 1.**
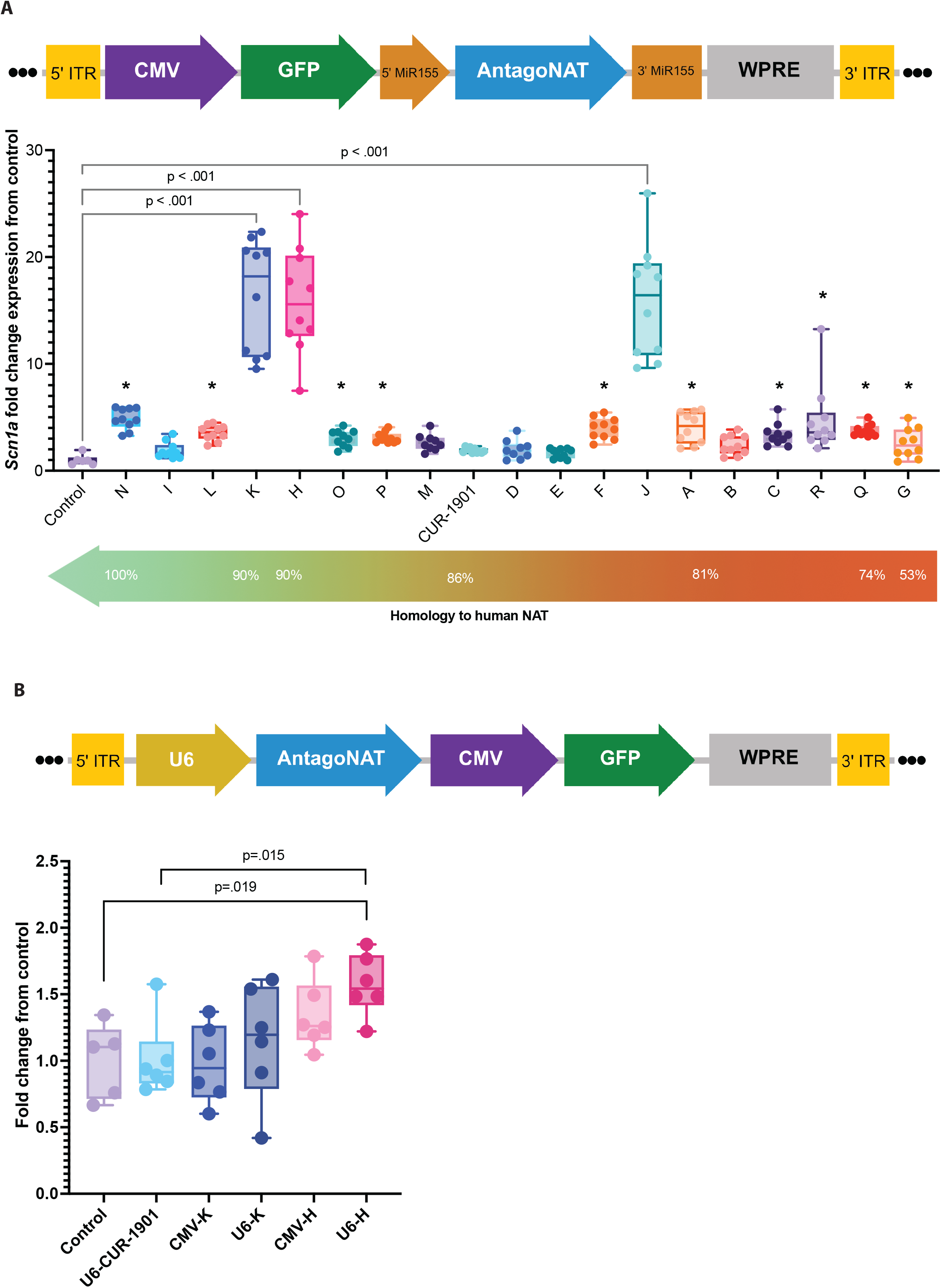
In vitro assessment of novel AntagoNAT sequences. (A) Schematic of AAV-CMV-eGFP-miR-155-AntagoNAT-miR-155-WPRE plasmid. Transient transfection revealed thirteen sequences to significantly increase *Scn1a* expression compared to the control cells (^*^ p<0.001). Candidates K, H and J showed the highest increase in expression (16-fold compared to control). One-Way ANOVA with multiple comparison correction using Two-stage linear step-up procedure of Benjamini, Krieger and Yekutieli. n=9/10 for AntagoNAT sequences, n=6 for control (3 technical replicates each). (B) Schematic of AAV-U6-AntagoNAT-CMV-eGFP-WPRE plasmid. In vitro transduction of AAV9 vectors (n=5/6 replicates of each group, 3 technical replicates each) using either the U6 or the CMV promoter, represented in a Box plot. One-way ANOVA, Dunnett’s multiple comparison performed.

### *In vitro* cell assays

30,000 Neuro2a cells were seeded per well in 24-well plates using differentiation media: DMEM (Thermo Fisher Scientific) supplemented with 2% FCS (Sigma-Aldrich), 0.5mM cAMP (Merck) and 20μM Retinoic Acid (Merck). At day 6, media was removed, and cells were transfected with plasmid (2μg) containing candidate AntagoNAT sequence using Lipofectamine 2000 (Thermo Fisher Scientific) in OptiMEM (Thermo Fisher Scientific) overnight. The next day the transfection media was removed and replaced with new differentiation media. For transduction experiments, media was removed from the cells and replaced with OptiMEM with a multiplicity of infection (MOI) of 1×10^6^ vector genomes per seeded cell. The cells were maintained for a further 5 days, at which point they were collected using TRIzol reagent (Thermo Fisher Scientific). RNA was extracted as described below; gene expression was determined by qPCR (see below).

### AAV vector production

Recombinant AAV vector was produced using a protocol previously described.^36^ AAVpro 293T cells (Cat. No. 632273, Takara Bio) were transfected with AAV plasmid carrying the AntagoNAT sequence, a AAV2 Rep & AAV9 Cap gene plasmid (University of Pennsylvania, USA) and adenovirus helper plasmid (Harvard University, USA) using polyethylenimine (PEImax, Polysciences Inc). The vector was purified in an AKTAprime plus HPLC machine (AKTA prime) using POROS CaptureSelect AAVX Resin (Thermo Fisher Scientific). The resulting AAV was treated with DNase I, and then titrated by qPCR (see below). Vector stock was normalised to 1×10^13^ vector genomes/ml.

### Animals

All procedures were performed in accordance with the UK Home Office Animals (Scientific Procedures) Act 1986. All mice were housed under a non-reversed 12 hour:12 hour light–dark cycle and had access to food and water *ad libitum*. Animals were housed in individually ventilated cages with access to environmental enrichment. 129S-*Scn1a*^tm1Kea^/Mmjax DS mouse strain which contain an exon 1 deletion of *Scn1a*, resulting in haploinsufficiency, were obtained from the Jackson laboratory (Jackson Laboratory, Maine, USA). The heterozygous mice were crossed with wild-type C57BL/6J mice (Charles River, UK). The resulting first generation (F1) of heterozygous mice was used for treatment assessment (F1; 129Sv x C57BL/6J, from here onward, called *Scn1a*^*+/-*^ in the text).^18,23^

### Neonatal and P14 injections

At post-natal day 0/1 (P0/1) pups received combination bilateral ICV and IV delivery. Bilateral ICV administration was performed where 5μl of AAV9 (5×10^9^ vg per hemisphere) or Dulbeccos Phosphate Buffered Saline (PBS, Thermo Fisher Scientific) was administered to each hemisphere using a 33-gauge hamilton needle (VWR), following the previously described coordinates.^37^ Subsequently, IV injection was performed by delivering 25μl of AAV9 vector (2.5×10^10^ vg per pup) or PBS to the superficial temporal vein using a 33-gauge needle.^38^ P14 mice were anesthetised with isoflurane and received bilateral ICV and IV injections. Bilateral ICV was performed where 10μl of AAV9 (5×10^9^ or 2.5×10^9^ vg per hemisphere) or PBS following previous protocols.^18^ Tail vein injections were performed, delivering 30μl of AAV9 vector (2.5×10^10^ or 5×10^9^vg vg per pup) or PBS. After the procedure, the mice were returned to their dam. Weight and survival were monitored.

### Temperature-induced seizures

Temperature-induced seizure assessment was conducted on *Scn1a*^*+/-*^ mice at P20. The mice were placed in a heated chamber and video recorded. Starting temperature was 37ºC, with incremental increases of 0.5ºC every 1 minute, until 43.5ºC. Seizures were recorded at the corresponding temperature and assigned a Racine score.^39^

### EEG recordings

DS mice (weight >20 grams) were anaesthetised with isoflurane (induction 5%, surgery 1.5%) and placed in a stereotaxic frame (Kopf). They were injected subcutaneously with buprenorphine (0.03 mg/ml) and Metacam (0.15 mg/ml). A wireless electrocorticogram (ECoG) transmitter (Cat. No. A3048P2-AA-C37-D, single-channel transmitter, Open Source Instruments) was implanted subcutaneously and the electrode was placed over the somatosensory cortex. EEG signals (sampled at 256 Hz) were recorded for 15 consecutive days for each animal. Epileptiform activity was analysed *post hoc*, while blinded to the treatment group, using Pyecog software. EEG recordings were manually assessed, defined as a pattern of repetitive spike discharges followed by a progressive evolution in spike amplitude with a distinct post-ictal depression phase.

### Tissue collection and stereoscopic microscopy

Mice were anaesthetised with isoflurane, an incision in the right atrium was made followed by perfusion by PBS to the left ventricle. Tissues were split in half, either stored in 4% PFA for 48 hours, and then 30% sucrose at 4ºC for immunohistochemistry, or snap frozen to -80ºC for molecular analysis. Analysis of GFP expression using a stereoscopic fluorescence microscope (MZ16F; Leica, Wetzlar, Germany). Representative images were captured using a digital microscope camera (DFC420; Leica Microsystems, Milton Keynes, UK) and software (Image Analysis; Leica Microsystems).^40^

### Immunohistochemistry

Brain samples for immunohistochemistry were sectioned to 40µm coronal sections using a sliding microtome (Carl Zeiss, Welwyn Garden City, UK), and stored at 4ºC in a solution of 15% sucrose in TBS, 30% ethylene glycol and 0.3% sodium azide. To visualise GFP immunoreactivity, immunohistochemistry was performed following previously published protocols.^41^

### DNA and RNA isolation

DNA extraction for tissues was carried out using the DNeasy Blood & Tissue kit (Qiagen) following manufacturer instructions. RNA extraction was performed using TRIzol reagent (Invitrogen) combined with the PureLink™ RNA Mini Kit (Invitrogen). For cells, media was removed, cells were washed with PBS, then 1ml of TRIzol per 1×10^5^ to 1×10^7^ cells was applied. The lysate was triturated several times and transferred into a 1.5ml tube. For snap frozen tissues, 1ml of TRIzol was added per 50-100mg. A 3mm stainless steel bead (Qiagen) was added and the tubes were placed on a TissueLyser II (Qiagen, UK) and homogenised at 30Hz for 3 minutes. The samples were processed immediately following the PureLink™ RNA Mini Kit instructions. Concentration was measured in a FLUOstar® Omega microplate reader (BMG Labtech). Reverse-transcription was performed using the High-Capacity cDNA Reverse Transcription kit (Applied Biosystems). The resulting cDNA was used for quantitative PCR (qPCR).

### qPCR

Primers and probes were designed to target the *Scn1a* and *Gapdh* genes. *Gapdh* was used as housekeeping. For *Scn1a;* forward: TCAGAGGGAAGCACAGTAGAC, reverse: TTCCACGCTGATTTGACAGCA, probe: CCAGAAGAAACCCTTGAGCCCGAA (fluorophore: ABY, quencher: QSY, sourced from Thermo Fisher Scientific). For Gapdh; forward: ACGGCAAATTCAACGGCAC, reverse: TAGTGGGGTCTCGCTCCTGG, probe: TTGTCATCAACGGGAAGCCCATCA (fluorophore: VIC, quencher: QSY, sourced from Thermo Fisher Scientific). AAV vector titration was performed using primers targeting GFP; forward: GGCACAAGCTGGAGTACAAC, reverse: AGTTCACCTTGATGCCGTTC, probe: AGCCACAACGTCTATATCATGGCCG (fluorophore: FAM, quencher: ZEN / Iowa Black™ FQ, sourced from Integrated DNA Technologies, IDT). Standards for all genes of interest were sourced as gBlocks from IDT and a standard curve from 10^9^ to 10^3^ copies were used in all experiments. For titrations, the DNase I treated vector was serially diluted 6 times and the average concentration from all dilutions within the standard curve was used as the vector titre. Luna® Universal Probe qPCR Master Mix (NEB) with 250nM of primers and probes was used in all reactions. 5µL of each cDNA/vector sample was used. The plate was run in a QuantStudio 3 instrument (Applied Biosystems). Data was analysed using the QuantStudio Design and Analysis v1.4 (Applied Biosystems) software. Technical replicates were assessed and accepted if they were between 0.5 CTs from each other.

### Capillary immunoassay

Tissues were homogenised in T-PER™ Tissue Protein Extraction Reagent (Thermo Fisher Scientific) with 1X cOmplete™, EDTA-free Protease Inhibitor Cocktail (Merck) using a Qiagen TissueLyser II (30 Hz for up to 2 minutes). Lysates were incubated on ice for 5 minutes then sonicated for a further 5 minutes. Lysates were centrifuged at 14,000xg at 4°C for 15 minutes after which supernatants containing the membrane-enriched fraction were centrifuged for a further 5 minutes. Concentrations of resulting protein lysates were determined using the BioRad DC protein assay (BioRad) according to manufacturer’s instructions. A capillary immunoassay using these protein lysates was performed on the ProteinSimple Jess System (Bio-techne) using the manufacurer’s template for a 66-440kDa RePlex chemiluminescence assay (Bio-techne) with Total Protein normalisation. 0.5mg/ml lysates and anti-Na_V_1.1 antibody (Alomone ASC-001; 1:50) were used as previously described.^42^ Automatic normalisation to Total Protein and quantification of Nav1.1 protein signal was achieved using Compass for Simple Western software (Bio-techne).

### Statistical analysis

GraphPad Prism (version 10.2.2, Boston, Massachusetts, USA) was used for statistical analysis. For *in vitro* data, One-Way ANOVA with multiple comparison, with a post-hoc analysis using the two-stage linear step-up procedure of Benjamini, Krieger and Yekutieli multiple comparison. For smaller group number comparisons, we used the post-hoc analysis using the Dunnett’s multiple comparison test. For *in vivo* assessment, the following statistical analysis were used: survival data was analysed using the Log-rank (Mantel-Cox), for weight Two-Way ANOVA with a post-hoc analysis using the Dunnett’s multiple comparisons test was performed. For EEG data, Two-Way ANOVA with a post-hoc analysis using the Greenhouse-Geisser correction was performed. To compare means between two groups from EEG data, Unpaired Mann-Whitney test was used. To assess DNA and RNA gene expression data we used One-Way ANOVA with a post-hoc analysis using the Šídák’s multiple comparisons test. Data presented as mean ± standard error mean (SEM). The *n* number for each experimental group is clarified in the figures and figure legends. *p*-values are also shown in figures. Normal distribution of the data was assessed with the D’Agostino-Pearson test.

### Data availability

The data that support the findings of this study are available from the corresponding author, upon request.

## Results

### Development of novel AntagoNAT sequences

To investigate whether additional AntagoNAT sequences provided superior survival and protection against seizures, compared to the published sequence (CUR-1901),^33^ we designed eighteen novel AntagoNAT sequences. We incorporated these into an AAV plasmid backbone, driven by a RNA polymerase II CMV promoter, driving the expression of enhanced GFP, followed by the AntagoNAT sequences, which were flanked by miR-155 sequences (Fig.1A).^35^ miR-155 sequences are short hairpin looped structures which allow stable transcription of small RNA sequences by RNA polymerase II promoters.^35^

We transfected the AAV backbone constructs into differentiated Neuro2a (N2a) cells. Thirteen of the novel AntagoNAT sequences significantly increased endogenous *Scn1a* expression compared to control group, ranging between 3.4±0.33-fold (candidate C) to 16±1.6-fold (Candidate H) (Fig. 1A). AntagoNAT sequences H, J and K showed the highest increase in *Scn1a* expression (∼16-fold increase each; *p*<0.001), with K and H being the most homologous to the human NAT sequence (90%). Candidate N also showed a significant increase in *Scn1a* and had a higher similarity to human NAT. However, in this study we aimed to obtain a significant increase of endogenous *Scn1a* expression *in vivo* coupled with a lower vector dose of administration and therefore, AntagoNAT-K and H were chosen for the *in vivo* pre-clinical study. CUR-1901,^33^ showed an increase in endogenous *Scn1a* expression to 1.9±0.07-fold compared to control, when delivered from our plasmid construct (Fig. 1A).

Before we progressed to the *in vivo* study, we compared the CMV promoter versus a U6 RNA polymerase III promoter. In general, RNA polymerase III promoters are suitable to drive small RNA sequences which do not require protein translation.^43^ AAV9 vectors were produced with both promoters and used to transduce N2a cells. The U6 promoter revealed a significant increase in endogenous *Scn1a* expression with AntagoNAT-H and an increased expression with AntagoNAT-K compared to control group (Fig. 1B), we therefore proceeded with the AAV9-U6-AntagoNAT-H and K vectors for the *in vivo* study.

### Neonatal delivery of novel AAV9-AntagoNAT-H reduces SUDEP and seizure phenotype in a Dravet syndrome mouse model

To test the efficacy of our novel AntagoNAT sequences, we produced an AAV9 vector to deliver AntagoNAT-H and K. We tested the effects of our AAV9 therapy in the *Scn1a*^+/−^ mice.^21^ We administered the AAV9-AntagoNAT-H & K to newborn *Scn1a*^*+/-*^ mice via bilateral intracerebroventricular (ICV) and intravenous (IV) delivery (3.5×10^10^ vg/mouse) in a blinded and randomised study (Fig. 2A). AAV9-AntagoNAT-H via ICV and IV significantly increased survival to 84.2% (*p*=0.04, Fig. 2B) compared to 50% of PBS control *Scn1a*^*+/-*^ mice at P100. No difference in weight was observed (Fig. 2C). In contrast, delivery of AAV9-AntagoNAT-K via ICV and IV revealed 58% survival compared to 50% of PBS control *Scn1a*^*+/-*^ mice at P100 (Supplementary Fig. 1). Thus, AAV9-AntagoNAT-H was our lead candidate for the rest of the pre-clinical gene therapy study.

**Figure 2.**
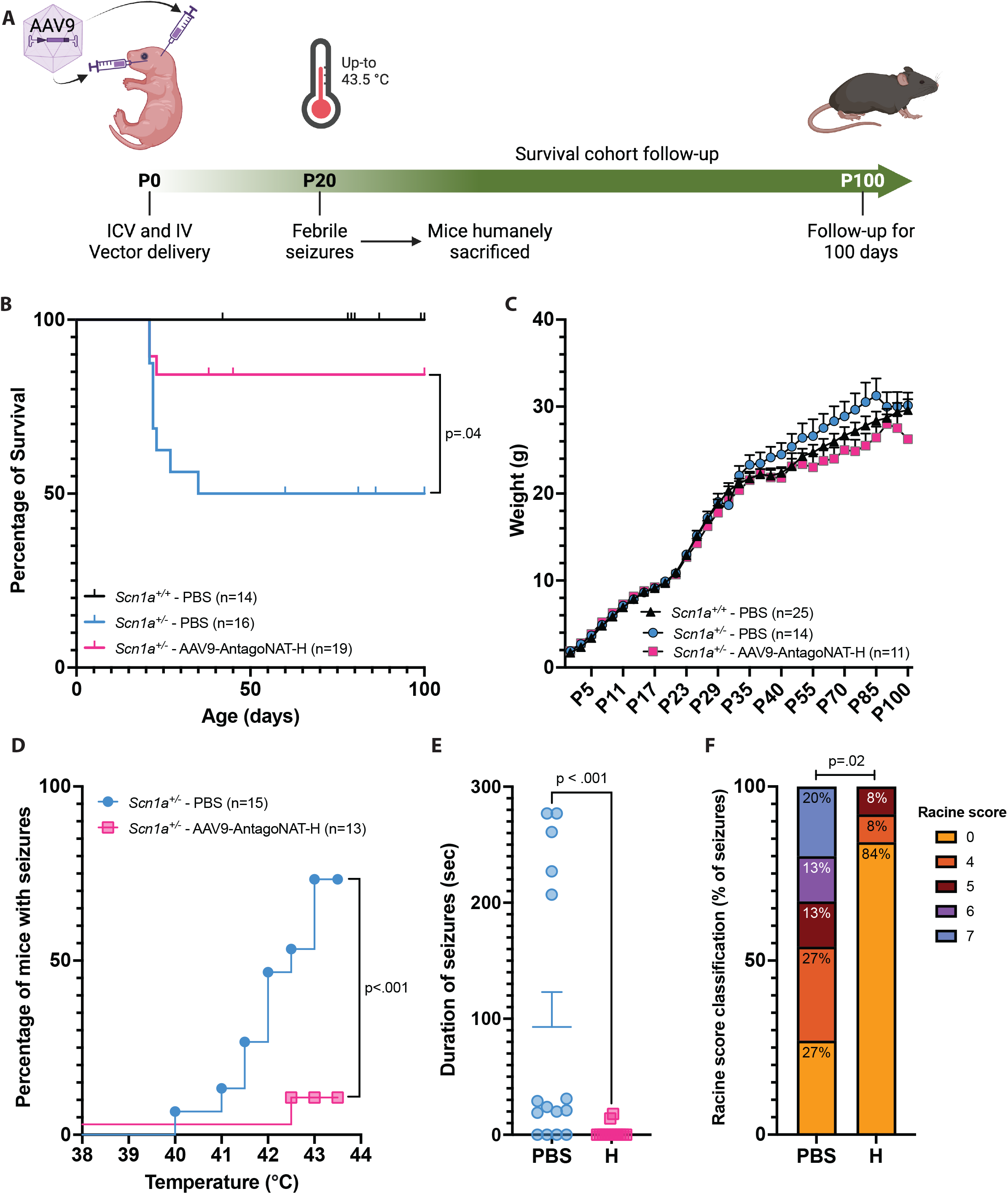
Neonatal AAV9-AntagoNAT-H gene therapy increases survival and reduces febrile seizures in DS mice. Scn1a^+/-^ mice received either AAV9-AntagoNAT-H or PBS via ICV and IV. PBS was also administered to wild type mice (Scn1a^+/+^). (A) Experiment timeline. (B) Survival of Scn1a^+/-^ DS mice treated with AAV9-AntagoNAT-H, shown in percentage of survival. Log-rank (Mantel-Cox) test. (C) Weight curves. Two-Way ANOVA with Dunnett’s multiple comparisons test. (D) Febrile seizure temperature threshold assessment. Log-rank (Mantel-Cox) test. (E) Duration of febrile seizures. Mann Whitney test. (F) The Racine score classification of febrile seizures. Fisher’s exact test.

We next examined the susceptibility to febrile seizures, as this is a common symptom amongst DS patients.^44^ By P25 we have observed approximately 40% of *Scn1a*^*+/-*^ mice undergo SUDEP, with mortality starting at P21 (Fig. 2B), therefore we conducted temperature-induced seizures just before this at P20. *Scn1a*^*+/-*^ mice which received ICV and IV AAV9-AntagoNAT-H or PBS at P0 were subjected to an increase of temperature from 37-43.5°C in increments of 0.5°C per minute. Gene therapy significantly reduced sensitivity to febrile seizures compared to PBS control mice, where 1 out of 13 treated mice exhibited a seizure and 11 out of 15 PBS control mice had seizures (*p*<0.001; Fig. 2D). Furthermore, seizure duration was significantly reduced (*p*<0.001; Fig. 2E). AAV9-AntagoNAT-H reduced mean Racine score (0.69±0.47 vs. PBS control 3.9±0.69; *p*<0.02) (Fig. 2F).

To assess if our AAV9-AntagoNAT-H treatment was effective in reducing spontaneous seizures, we measured seizure frequency via EEG recordings for 15 days (P30-45, Fig. 3A). We observed a significant reduction (*p*=0.025; Fig. 3B) in number of seizures over 15 days in gene therapy treated *Scn1a*^*+/-*^ mice compared to PBS control mice. The average daily seizures per mouse for AAV9-AntagoNAT-H group were 0.03±0.02 compared to 1.1±0.43 for PBS control group (Fig. 3C), We also observed a decrease in seizure duration between treatment groups over 15 days, from 25.14±9.32 to 2.56±1.67 seconds (Fig. 3D). Overall, in this neonatal study we observed a substantial increase in survival and a decrease in seizures by both febrile and EEG seizure recordings.

**Figure 3.**
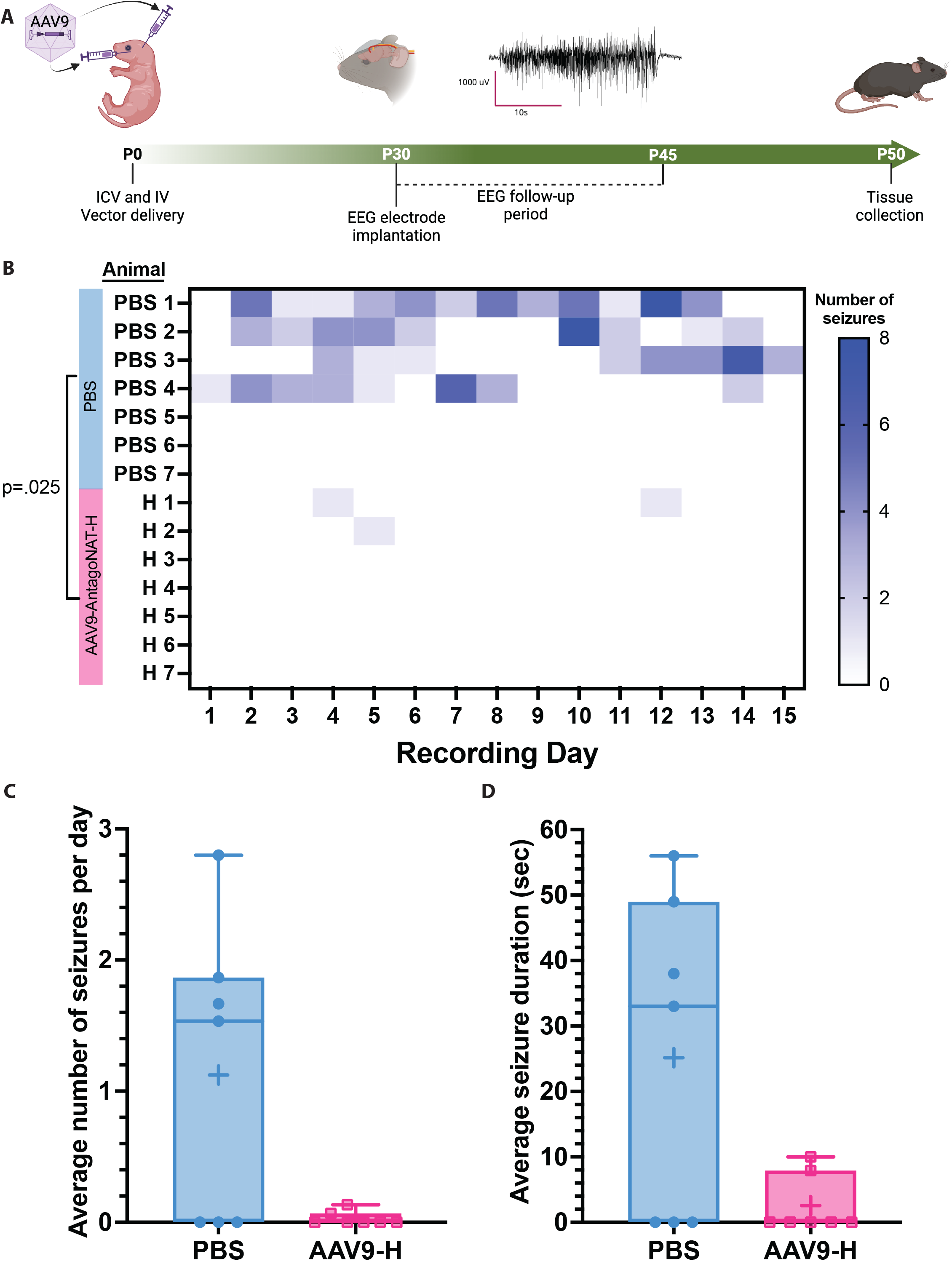
Improved seizure phenotype after neonatal gene transfer of AAV9-AntagoNAT-H. (A) Experiment schematic. (B) Heat map illustrating seizure events per day over 15 days. Two-Way ANOVA with a post-hoc analysis using the Greenhouse-Geisser correction. (C) The average number of seizure events per mouse per day. Unpaired Mann-Whitney test. (D) Average seizure duration over the recording period between the AAV9-AntagoNAT-H and PBS control groups. Unpaired Mann-Whitney test.

*Scn1a* mRNA expression was measured by qPCR of cerebral cortex and heart samples of P20 mice. In the cerebral cortex we observed a significant increase in endogenous *Scn1a* mRNA in AAV9-AntagoNAT-H *Scn1a*^*+/-*^ group, compared to PBS group (*p*=0.036; Fig. 4A). In the heart, we observed an increased trend of endogenous *Scn1a* mRNA in treated group compared to PBS controls (Fig. 4B). We also assessed the number of vector copies present in both the cerebral cortex and the heart. As expected, the vector copies were significantly higher in AAV9-AntagoNAT-H *Scn1a*^*+/-*^ group compared to control groups (cerebral cortex; *p*<0.001, heart: *p*=0.004; Fig. 4C and D). Furthermore, Na_V_1.1 expression in the cerebral cortex also showed an increased trend in AAV9-AntagoNAT-H *Scn1a*^*+/-*^ group (0.49±0.06) but this was not significant when compared to PBS control group (0.44±0.03) (Fig. 4E).

**Figure 4.**
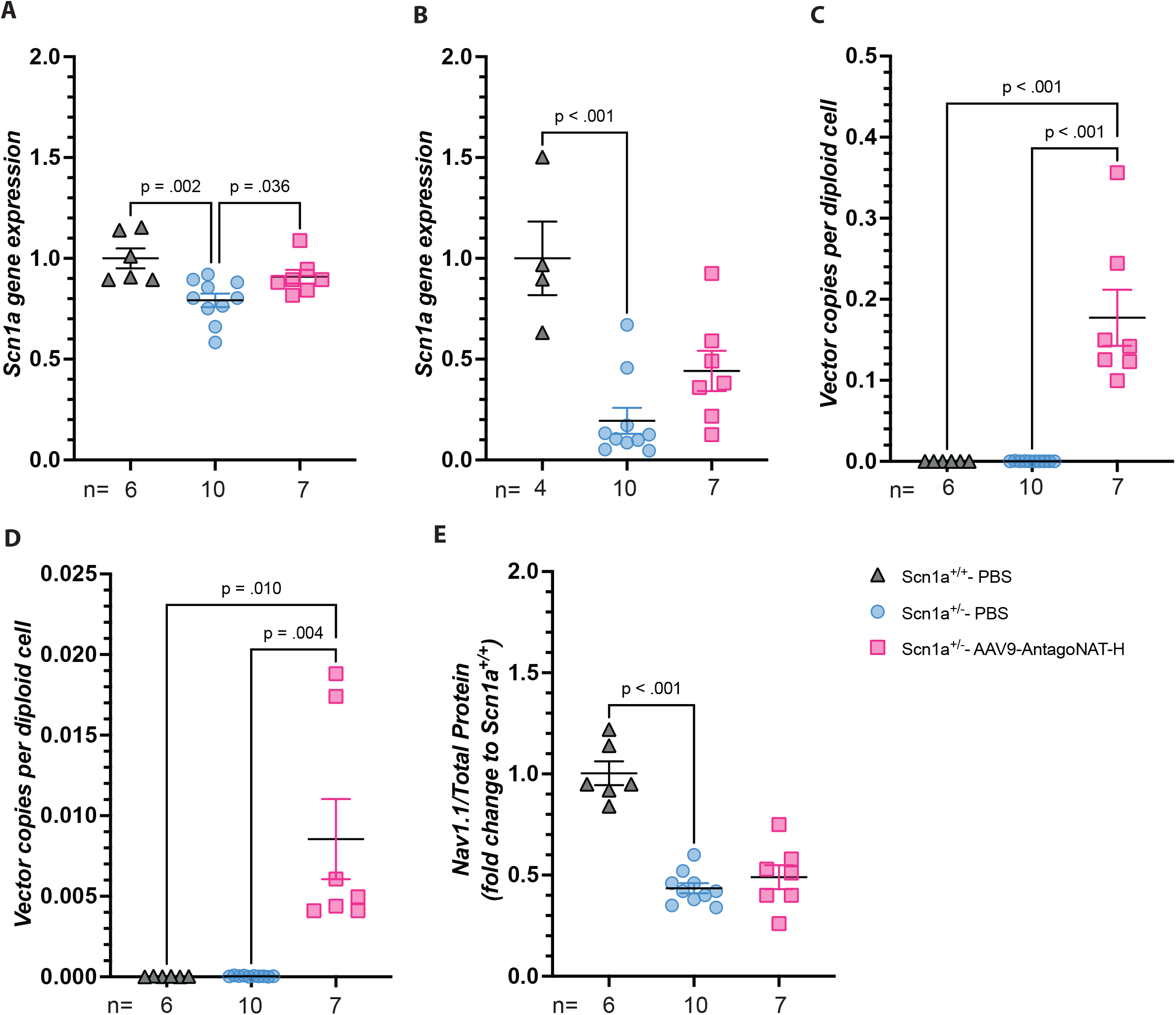
Increase of endogenous Scn1a and Na_V_1.1 in the cortex and heart of AAV9-AntagoNAT-H treated Scn1a^+/-^ DS mice. (A) *Scn1a* expression in the cortex. (B) *Scn1a* expression in the heart. (C) Vector copy number analysis in the cortex. (D) Vector copy number in the heart. (E) Na_V_1.1 expression in the cortex. One-Way ANOVA Holm-Šídák’s multiple comparisons test. n numbers for each group are indicated in the figure.

### P14 delivery of novel AAV9-AntagoNAT-H reduces SUDEP

The *Scn1a* transcript becomes stable at P14 of development.^18^ We therefore asked whether delivering our AAV9-AntagoNAT-H treatment at P14, closer to the time point to seizure and SUDEP onset, could prolong survival in *Scn1a*^*+/-*^ DS model. We thus delivered AAV9-AntagoNAT-H to P14 *Scn1a*^*+/-*^ mice in a blinded and randomised study (Fig. 5A). P14 *Scn1a*^*+/-*^ mice initially received the same dose as in the neonatal study (ICV; 5×10^9^vg per hemisphere and IV; 2.5×10^10^vg, for a total dose of 3.5×10^10^ vg/mouse). This dose was not efficacious, as the treated AAV9-AntagoNAT-H *Scn1a*^*+/-*^ mice revealed an overall survival of 14% compared to 69% of PBS control mice (Supplementary Fig. 2).

**Figure 5.**
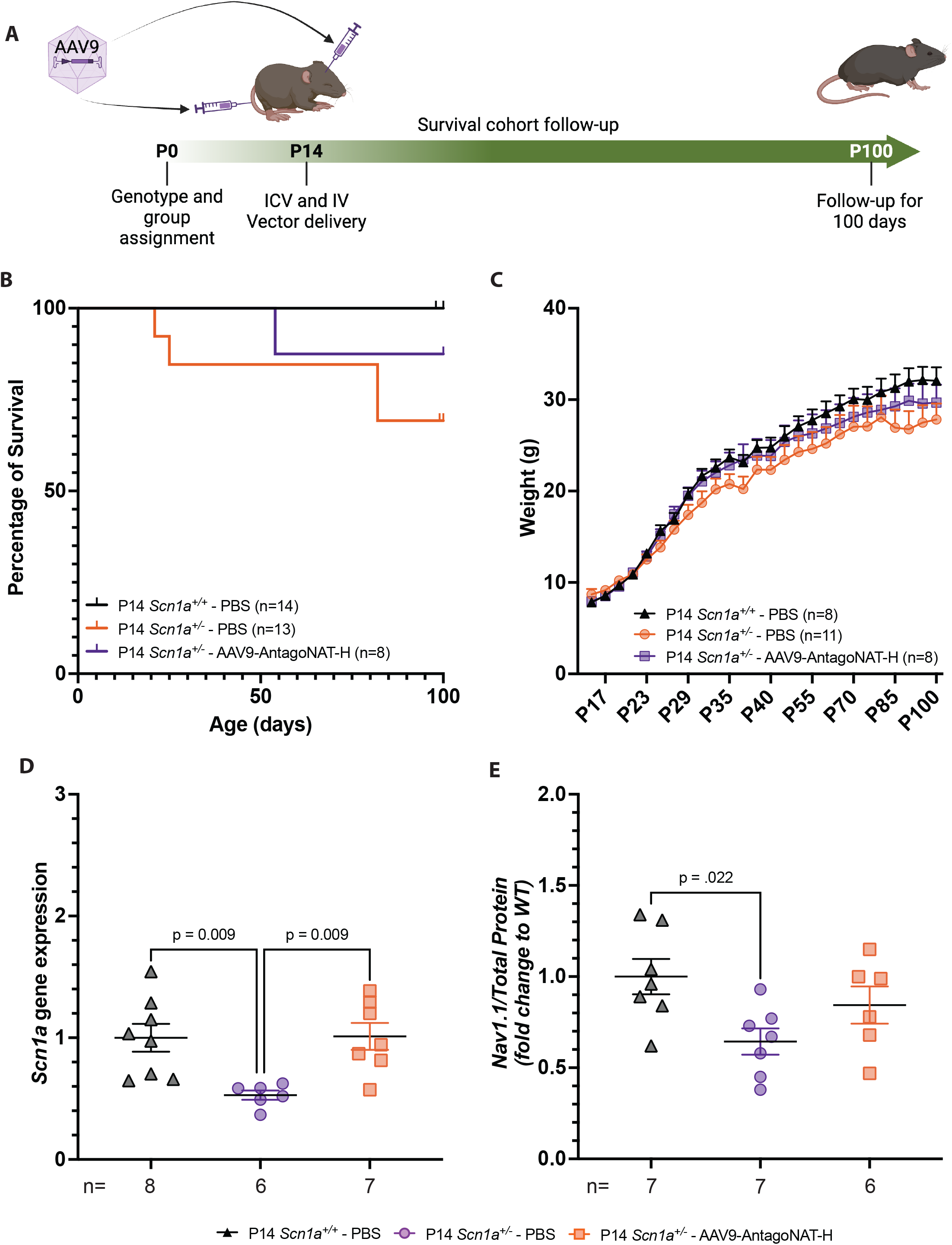
Reduction in SUDEP after ICV and IV AAV9-AntagoNAT-H therapy to P14 Scn1a+/-DS mice. (A) Schematic diagram showing the experimental plan. (B) Survival curve, showing percentage of survival. Log-rank (Mantel-Cox) test. (C) Weights. Two-Way ANOVA with Greenhouse-Geisser correction. (D) *Scn1a* expression in cerebral cortex. Cerebral cortex tissues for Scn1a^+/-^ group, range from P20-100. E) Na_V_1.1 expression in the cerebral cortex. One-Way ANOVA Holm-Šídák’s multiple comparisons test.

We, therefore, administered a lower dose. The P14 *Scn1a*^*+/-*^ mice received 2.5×10^9^ vg per hemisphere, via bilateral ICV and 5×10^9^ vg via tail vein injection (total dose of 1×10^10^ vg/mouse) and were monitored up to P100. We observed an 87.5% survival in our AAV9-AntagoNAT-H *Scn1a*^*+/-*^ group compared to 69% of PBS control mice (Fig. 5B). No difference in weight was observed (Fig. 5C). *Scn1a* mRNA expression was measured by qPCR of cerebral cortex and we observed a significant increase in endogenous *Scn1a* in treated group compared to PBS control group (Fig. 5D). We were unable to detect *Scn1a* in the heart. Furthermore, AAV9-AntagoNAT-H *Scn1a*^*+/-*^ group showed an increased trend of Na_V_1.1 expression in the cerebral cortex compared to PBS control group (Fig. 5E).

## Discussion

The novel AAV9-AntagoNAT-H therapy delivered to neonatal *Scn1a*^*+/-*^ mice via combination administration (ICV and IV) significantly improves the disease phenotype in the *Scn1a*^*+/-*^ DS mice; reduces SUDEP (Fig. 2B), susceptibility to febrile seizures (Fig. 2D-F), and seizure frequencies as assessed via EEG recordings (Fig. 3B-D). In addition, we measured a significant increase in endogenous *Scn1a* mRNA and observed an increased trend of Na_V_1.1 protein in the cortex (Fig. 4A and E) of our treatment group compared to controls. We also observed an increase of endogenous *Scn1a* in the hearts of treated *Scn1a*^*+/-*^ mice (Fig. 4B and E). AAV9-AntagoNAT-H was administered to juvenile *Scn1a*^*+/-*^ mice (P14), at a time point when Na_V_1.1 expression is stable and this revealed an increase in survival. Thus, we demonstrated an improvement on previous publication, where AntagoNATs were delivered via repeated intrathecal delivery to DS mice, ^33^ by designing new sequences and incorporating them into an AAV9 viral vector and therefore providing a one-off treatment. More importantly, due to the broad biodistribution of AAV9 viral vectors^45^ and mechanism of action of the AntagoNATs,^33^ which should only have an effect in cells which express the *Scn1a* mRNA,^33^ we have validated a therapy which is able to achieve an extensive transduction profile after single administration (via ICV and IV) and potentially target all cells expressing *Scn1a*. Therefore, we have not restricted our gene therapy to GABAergic inhibitory interneurons, which has been previously demonstrated in other gene therapy and editing pre-clinical studies for DS.^19,46^

In this study, we aimed to target the brain and heart by dual administration (bilateral ICV and IV) of AAV9-AntagoNAT-H therapy. As previous studies have detected Na_V_1.1 expression in inhibitory interneurons,^7^ Purkinje neurons,^11^ CA1 pyramidal cells^12^ and the heart.^16^ In the neonatal study we achieved a significant improvement in survival and a reduction in SUDEP events in our AAV9-AntagoNAT-H treated group. Further evidence of efficacy was observed in the assessment of febrile seizures, which are also observed in DS patients.^1^ The AAV9-AntagoNAT-H treated group showed a significant reduction of seizure events and a reduction in severity compared to PBS controls. Spontaneous seizure events were assessed by EEG recordings from P30-45 of development. We observed a reduction in seizure frequency in the AAV9-AntagoNAT-H treated *Scn1a*^*+/-*^ mice compared to PBS controls. Our results corroborate with previous pre-clinical studies.^18,19^Although these results are encouraging, further studies are required to increase the statistical power of this outcome measure.

In the neonatal study we demonstrated a significant increase in endogenous *Scn1*a mRNA expression in the cerebral cortex and an increased trend of expression in the heart. Interestingly, we did not achieve a significant increase in Na_V_1.1 expression in the cerebral cortex of treated mice and we were unable to detect Na_V_1.1 in the heart. This may be due to the known difficulties with Na_V_1.1 antibodies which previous studies have addressed by using reporter tags surrounding the *SCN1A* gene, to identify the Na_V_1.1 expression in DS mice.^30,47^ Also low levels of Na_V_1.1 have been reported in human and dog hearts^16^ and therefore, the Na_V_1.1 levels in a mouse heart perhaps too low to detect. Future studies, would require assessment of sodium current specifically from Na_V_1.1 in cardiomyocytes.^16^ The Na_V_1.1 protein results in the cerebral cortex contrast with previous data, where a significant increase of Na_V_1.1 was detected in the brain after a single neonatal ICV administration of AAV9^19^ or ASO^18^ therapy to *Scn1a*^*+/-*^ DS mice. Our data could also provide evidence that normalisation of in Na_V_1.1 expression to wild-type levels may not be entirely required to have a therapeutic benefit.

In this study, we aimed to deliver our novel AAV9-AntagoNAT-H therapy to older P14 *Scn1a*^*+/-*^ DS mice as the *Scn1a* transcript in the brain is stable at this time point.^18^ In addition, the rodent brain development at P14 is comparable to brain development in a 1-2 year old human brain.^48^ Treatment at P14 did not reveal a significant increase in survival compared to control mice (Fig. 5B), However, this is the first study, to our knowledge, in which an AAV based therapy has been delivered to P14 *Scn1a*^*+/-*^ DS mice. Further studies using an alternative heterozygous mouse model, where there is a longer period between first seizure and SUDEP,^30^ would be useful to further validate our treatment and to address the efficiency in older *Scn1a*^*+/-*^ DS mice after disease symptom onset.

To further support the clinical translation of our AAV9-AntagoNAT-H gene therapy approach. A recent study examined the expression profile of *SCN1A* lncRNA in brain samples from drug resistant epilepsy patients.^49^ The study revealed that the *SCN1A* NAT is abundantly expressed across frontal and temporal lobe from brain samples between the ages of 1-19 years.^49^ This study validated the expression profile of *SCN1A* NAT in a human brain and therefore confirms the stable expression profile of our gene therapy target. Moreover, it supports and provides confidence for the applicability of the AAV9-AntagoNAT approach to be used in patients between the ages of 1-19 years.

There are a number of genetic therapeutic strategies developed for DS, including the AntagoNAT which has shown decrease in seizure frequency after repeated intrathecal administration in a DS knock-in model.^33^ Stoke Therapeutics have developed ASO therapy which has shown a reduction of convulsive seizures in a Phase 1/2a clinical trial after repeated intrathecal administration to DS patients between the ages of 2-18.^31^ Encoded Therapeutics have developed an AAV9 transcription factor approach targeting GABAergic interneurons, where the pre-clinical study showed reduction of seizure phenotype and restoration of *Scn1a* and Na_V_1.1 expression after single neonatal ICV delivery.^19^ Encoded Therapeutics have recently announced start date for Phase 1/2 clinical trials.^32^ Our AAV9-AntagoNAT therapy, like Encoded Therapeutics’ approach, provides a one-off treatment, however, the added advantage of AAV9-AntagoNAT is that it is able to target additional cells in the CNS which also express Na_V_1.1 protein. The addition of IV delivery of our AAV9-AntagoNAT therapy was to achieve efficient targeting of the heart, as the heart has shown to contribute to SUDEP through altered electrical function.^13,14^ However, we were unable to show sufficient increase of *Scn1a* expression and unable to detect Na_V_1.1 expression in the neonatal and P14 administration studies. Further studies, examining the electrical function in the heart after gene therapy would be useful to address the association with SUDEP.

Here, we provide proof of concept pre-clinical evidence that neonatal ICV and IV administration of AAV9-AntagoNAT-H, designed to specifically target *Scn1a* and *SCN1A* lncRNA can effectively restore *Scn1a* gene expression, modestly increase Na_V_1.1 production and reduces SUDEP incidences and seizures in a clinically relevant *Scn1a*^*+/-*^ DS mouse model. Additionally, administration at P14 showed a reduction in SUDEP and increase in endogenous *Scn1a* and Na_V_1.1 expression. In summary, the AAV9-AntagoNAT strategy, provides great promise as a genetic therapy for DS and requires further pre-clinical testing to evaluate the safety and efficiency of this therapy.

## Supporting information

Seizure-Treated mice

Seizure-untreated mice

## Acknowledgements

Figures 2A, 3A and 5A were created with BioRender.com

---

**Supplementary Figure 1.**
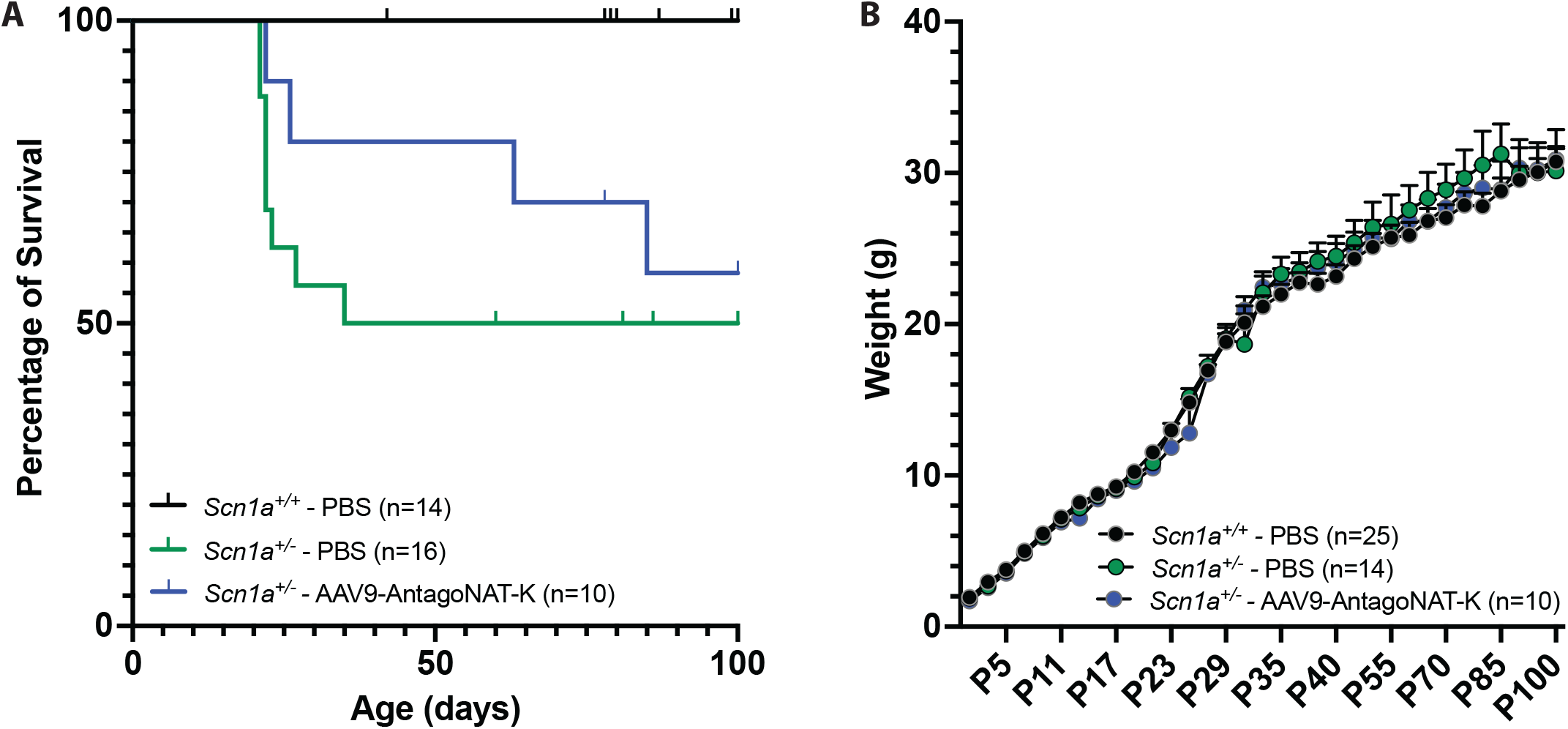
Survival of AAV9-AntagoNAT-K. (A) Survival curve of Scn1a^+/-^ DS mice treated with AAV9-AntagoNAT-K (*p*=0.418). Log-rank (Mantel-Cox) test. (B) Weights. Two-Way ANOVA with Dunnett’s multiple comparisons test.

**Supplementary Figure 3.**
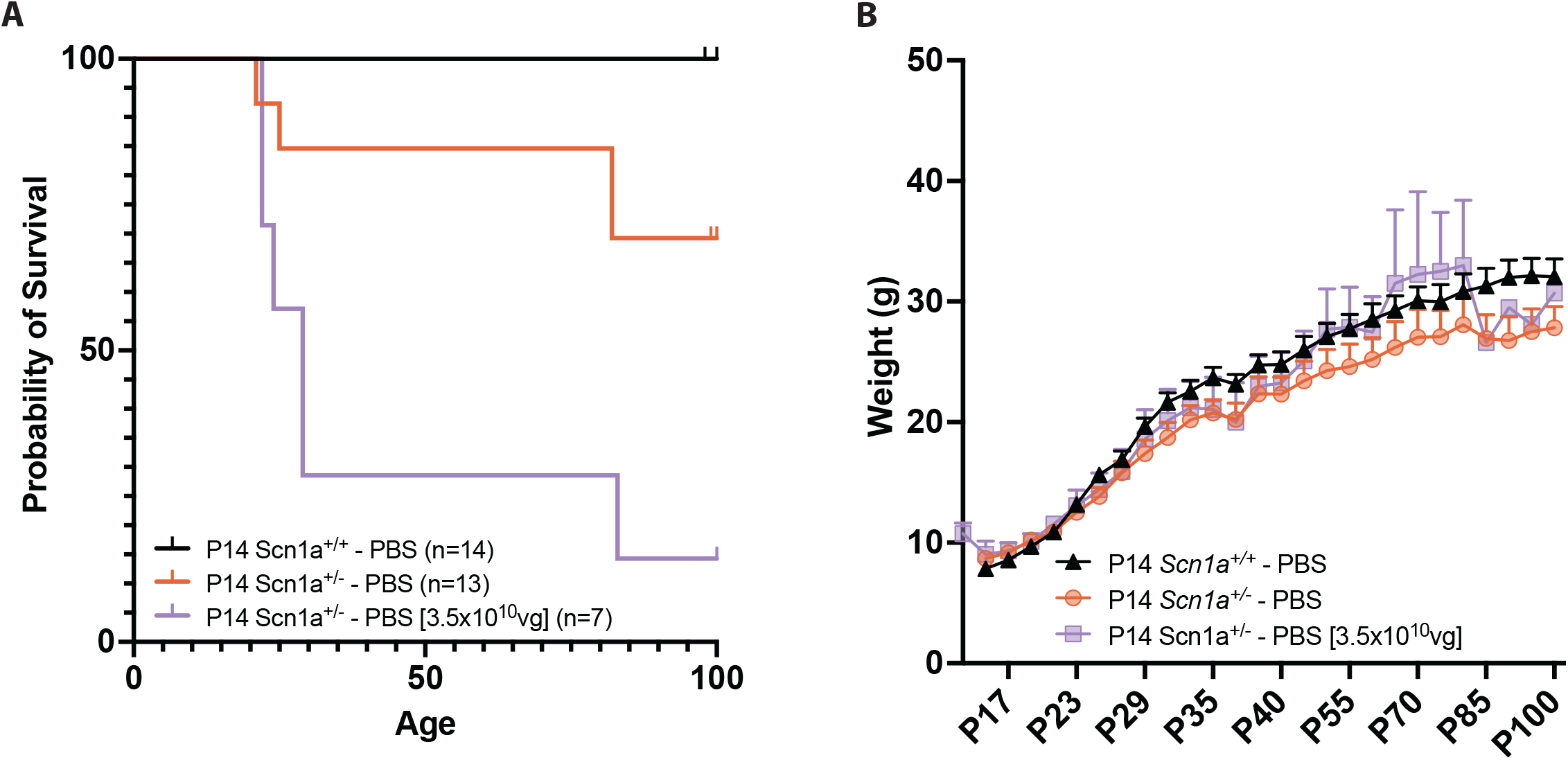
Survival of P14 Scn1a^+/-^ mice treated with AAV9-AntagoNAT-H (dose; 3.5×10^10^vg/mouse). (A) Survival curve of Scn1a^+/-^ DS mice treated with AAV9-AntagoNAT-H. Log-rank (Mantel-Cox) test. (B) Weights.Two-Way ANOVA with Dunnett’s multiple comparisons test.

## References

1. Chilcott E, Díaz JA, Bertram C, Berti M, Karda R. Genetic therapeutic advancements for Dravet Syndrome. Epilepsy & Behavior. 2022;132:108741.

2. Strzelczyk A, Lagae L, Wilmshurst J, et al. Dravet syndrome: a systematic literature review of the illness burden. Epilepsia Open. 2023: 1256–1270.

3. Brunklaus A, Zuberi SM. Dravet syndrome—From epileptic encephalopathy to channelopathy. Epilepsia. 2014;55(7):979–984.

4. Brunklaus A, Pérez-Palma E, Ghanty I, et al. Development and Validation of a Prediction Model for Early Diagnosis of SCN1A-Related Epilepsies. Neurology. 2022;98(11):e1163–e1174.

5. Wolff M, Cassé-Perrot C, Dravet C. Severe myoclonic epilepsy of infants (Dravet syndrome): natural history and neuropsychological findings. Epilepsia. 2006;45–48.

6. Whitaker WRJ, Faull RLM, Waldvogel HJ, Plumpton CJ, Emson PC, Clare JJ. Comparative distribution of voltage-gated sodium channel proteins in human brain. Molecular Brain Research. 2001;88(1):37–53.

7. Yu FH, Mantegazza M, Westenbroek RE, et al. Reduced sodium current in GABAergic interneurons in a mouse model of severe myoclonic epilepsy in infancy. Nature neuroscience. 2006;9(9):1142–1149.

8. Ogiwara I, Miyamoto H, Morita N, et al. Nav1.1 Localizes to Axons of Parvalbumin-Positive Inhibitory Interneurons: A Circuit Basis for Epileptic Seizures in Mice Carrying an Scn1a Gene Mutation. Journal of Neuroscience. 2007;27(22):5903–5914.

9. Schaller KL, Caldwell JH. Expression and distribution of voltage-gated sodium channels in the cerebellum. Cerebellum. 2003;2(1):2–9.

10. Beckh S, Noda M, Lübbert H, Numa S. Differential regulation of three sodium channel messenger RNAs in the rat central nervous system during development. EMBO J. 1989;8(12):3611–3616.

11. Kalume F, Frank HY, Westenbroek RE, Scheuer T, Catterall WA. Reduced sodium current in Purkinje neurons from Nav1. 1 mutant mice: implications for ataxia in severe myoclonic epilepsy in infancy. Journal of Neuroscience. 2007;27(41):11065–11074.

12. Almog Y, Mavashov A, Brusel M, Rubinstein M. Functional investigation of a neuronal microcircuit in the CA1 area of the hippocampus reveals synaptic dysfunction in Dravet syndrome mice. Frontiers in Molecular Neuroscience. 2022;15:823640.

13. Berg AT, Coffman K, Gaebler-Spira D. Dysautonomia and functional impairment in rare developmental and epileptic encephalopathies: the other nervous system. Developmental Medicine & Child Neurology. 2021;63(12):1433–1440.

14. Auerbach DS, Jones J, Clawson BC, et al. Altered Cardiac Electrophysiology and SUDEP in a Model of Dravet Syndrome. PLoS ONE. 2013;8(10):1-15. d

15. Kalume F, Westenbroek RE, Cheah CS, et al. Sudden unexpected death in a mouse model of Dravet syndrome. J Clin Invest. 2013;123(4):1798–1808.

16. Mishra S, Reznikov V, Maltsev VA, Undrovinas NA, Sabbah HN, Undrovinas A. Contribution of sodium channel neuronal isoform Nav1.1 to late sodium current in ventricular myocytes from failing hearts. J Physiol. 2015;593(Pt 6):1409–1427.

17. Miller AR, Hawkins NA, Mccollom CE, Kearney JA. Mapping genetic modifiers of survival in a mouse model of Dravet syndrome. Genes, Brain and Behavior. 2014;13(2):163–172.

18. Han Z, Chen C, Christiansen A, et al. Antisense Oligonucleotides Increase Scn1a Expression and Reduce Seizures and SUDEP Incidence in a Mouse Model of Dravet Syndrome. Science translational medicine. 2020:eaaz6100

19. Tanenhaus A, Stowe T, Young A, et al. Cell-Selective Adeno-Associated Virus-Mediated SCN1A Gene Regulation Therapy Rescues Mortality and Seizure Phenotypes in a Dravet Syndrome Mouse Model and Is Well Tolerated in Nonhuman Primates. Hum Gene Ther. 2022;33(11-12):579–597.

20. Gerbatin RR, Augusto J, Boutouil H, Reschke CR, Henshall DC. Sexual dimorphism in epilepsy and comorbidities in Dravet syndrome mice carrying a targeted deletion of exon 1 of the Scn1a gene. Published online August 28, 2021:2021.08.27.457904.

21. Mistry AM, Thompson CH, Miller AR, Vanoye CG, George AL, Kearney J a. Strain- and age-dependent hippocampal neuron sodium currents correlate with epilepsy severity in Dravet syndrome mice. Neurobiology of Disease. 2014;65:1–11.

22. Cheah CS, Yu FH, Westenbroek RE, et al. Specific deletion of NaV1.1 sodium channels in inhibitory interneurons causes seizures and premature death in a mouse model of Dravet syndrome. Proc Natl Acad Sci U S A. 2012;109(36):14646–14651.

23. Miller AR, Hawkins NA, McCollom CE, Kearney JA. Mapping genetic modifiers of survival in a mouse model of Dravet syndrome. Genes Brain Behav. 2014;13(2):163–172.

24. Jones SP, O’Neill N, Muggeo S, Colasante G, Kullmann DM, Lignani G. Developmental instability of CA1 pyramidal cells in Dravet Syndrome. bioRxiv. [Preprint] doi: 10.1101/2022.09.12.507264

25. Myers KA, Bello-Espinosa LE, Symonds JD, et al. Heart rate variability in epilepsy: A potential biomarker of sudden unexpected death in epilepsy risk. Epilepsia. 2018;59(7):1372–1380.

26. Cardenal-Muñoz E, Auvin S, Villanueva V, et al. Guidance on Dravet syndrome from infant to adult care: Road map for treatment planning in Europe. Epilepsia Open. 2022;7(1):11–26.

27. Mendell JR, Al-Zaidy S, Shell R, et al. Single-Dose Gene-Replacement Therapy for Spinal Muscular Atrophy. New England Journal of Medicine. 2017;377(18):1713–1722.

28. Strauss KA, Farrar MA, Muntoni F, et al. Onasemnogene abeparvovec for presymptomatic infants with two copies of SMN2 at risk for spinal muscular atrophy type 1: the Phase III SPR1NT trial. Nat Med. 2022;28(7):1381–1389.

29. Grieger JC, Samulski RJ. Packaging Capacity of Adeno-Associated Virus Serotypes: Impact of Larger Genomes on Infectivity and Postentry Steps. J Virol. 2005;79(15):9933–9944.

30. Fadila S, Beucher B, Dopeso-Reyes IG, et al. Viral vector–mediated expression of NaV1.1, after seizure onset, reduces epilepsy in mice with Dravet syndrome. J Clin Invest. 2023;133(12):e159316.

31. Stoke Therapeutics Announces Positive New Safety & Efficacy Data from Patients Treated with STK-001 in the Phase 1/2a Studies (MONARCH & ADMIRAL) and the SWALLOWTAIL Open-Label Extension (OLE) Study in Children and Adolescents with Dravet Syndrome - Stoke Therapeutics. Accessed August 11, 2023. https://investor.stoketherapeutics.com/news-releases/news-release-details/stoke-therapeutics-announces-positive-new-safety-efficacy-data/

32. Encoded Therapeutics Announces US IND Clearance and Australian CTA Approval for Dravet Syndrome Gene Therapy Candidate ETX101. Encoded Therapeutics, Inc. Accessed May 21, 2024. https://encoded.com/press-releases/encoded-therapeutics-announces-us-ind-clearance-and-australian-cta-approval-for-dravet-syndrome-gene-therapy-candidate-et×101/

33. Hsiao J, Yuan TY, Tsai MS, et al. Upregulation of Haploinsufficient Gene Expression in the Brain by Targeting a Long Non-coding RNA Improves Seizure Phenotype in a Model of Dravet Syndrome. EBioMedicine. 2016;9:257–277.

34. Zuker M. Mfold web server for nucleic acid folding and hybridization prediction. Nucleic Acids Res. 2003;31(13):3406–3415.

35. Chung KH, Hart CC, Al-Bassam S, et al. Polycistronic RNA polymerase II expression vectors for RNA interference based on BIC/miR-155. Nucleic Acids Res. 2006;34(7):e53.

36. Keshavan N, Greenwood M, Prunty H, et al. Gene Therapy Prevents Hepatic Mitochondrial Dysfunction in Murine Deoxyguanosine Kinase Deficiency. Published online May 14, 2024:2024.05.10.593325.

37. Kim JY, Ash RT, Ceballos-Diaz C, et al. Viral transduction of the neonatal brain delivers controllable genetic mosaicism for visualising and manipulating neuronal circuits in vivo. European Journal of Neuroscience. 2013;37(8):1203–1220.

38. Delhove JMKM, Buckley SMK, Perocheau DP, et al. Longitudinal in vivo bioimaging of hepatocyte transcription factor activity following cholestatic liver injury in mice. Scientific Reports. 2017;7(October 2016):41874.

39. Van Erum J, Van Dam D, De Deyn PP. PTZ-induced seizures in mice require a revised Racine scale. Epilepsy & Behavior. 2019;95:51–55.

40. Rahim AA, Wong AMS, Hoefer K, et al. Intravenous administration of AAV2/9 to the fetal and neonatal mouse leads to differential targeting of CNS cell types and extensive transduction of the nervous system. The FASEB Journal. 2011;25(10):3505–3518.

41. Karda R, Rahim AA, Wong AMS, et al. Generation of light-producing somatic-transgenic mice using adeno-associated virus vectors. Scientific Reports. 2020;10(1):1–9.

42. Mavashov A, Brusel M, Liu J, et al. Heat-induced seizures, premature mortality, and hyperactivity in a novel Scn1a nonsense model for Dravet syndrome. Front Cell Neurosci. 2023;17:1149391.

43. Gao Z, Herrera-Carrillo E, Berkhout B. RNA Polymerase II Activity of Type 3 Pol III Promoters. Mol Ther Nucleic Acids. 2018;12:135–145.

44. Wolff M, Cassé-Perrot C, Dravet C. Severe Myoclonic Epilepsy of Infants (Dravet Syndrome): Natural History and Neuropsychological Findings. Epilepsia. 2006;47(2):45-48.

45. Mattar CN, Wong AMS, Hoefer K, et al. Systemic gene delivery following intravenous administration of AAV9 to fetal and neonatal mice and late-gestation nonhuman primates. FASEB J. 2015;29(9):3876–3888.

46. Colasante G, Lignani G, Brusco S, et al. dCas9-based Scn1a gene activation restores inhibitory interneuron excitability and attenuates seizures in Dravet syndrome mice. Molecular Therapy. 2020;28(1):235–253.

47. Mich JK, Ryu J, Wei AD, et al. AAV-mediated interneuron-specific gene replacement for Dravet syndrome. bioRxiv. [Preprint] doi: 10.1101/2023.12.15.571820

48. Semple BD, Blomgren K, Gimlin K, Ferriero DM, Noble-Haeusslein LJ. Brain development in rodents and humans: Identifying benchmarks of maturation and vulnerability to injury across species. Prog Neurobiol. 2013;0:1-16. d

49. Schneider MF, Vogt M, Scheuermann J, et al. Brain expression profiles of two SCN1A antisense RNAs in children and adolescents with epilepsy. Transl Neurosci. 2024;15(1):20220330.

